# A dual role for shape skeletons in human vision: perceptual organization and object recognition

**DOI:** 10.1101/799650

**Authors:** Vladislav Ayzenberg, Frederik S. Kamps, Daniel D. Dilks, Stella F. Lourenco

## Abstract

Shape perception is crucial for object recognition. However, it remains unknown exactly how shape information is represented, and, consequently, used by the visual system. Here, we hypothesized that the visual system represents “shape skeletons” to *both* (1) perceptually organize contours and component parts into a shape percept, and (2) compare shapes to recognize objects. Using functional magnetic resonance imaging (fMRI) and representational similarity analysis (RSA), we found that a model of skeletal similarity explained significant unique variance in the response profiles of V3 and LO, regions known to be involved in perceptual organization and object recognition, respectively. Moreover, the skeletal model remained predictive in these regions even when controlling for other models of visual similarity that approximate low- to high-level visual features (i.e., Gabor-jet, GIST, HMAX, and AlexNet), and across different surface forms, a manipulation that altered object contours while preserving the underlying skeleton. Together, these findings shed light on the functional roles of shape skeletons in human vision, as well as the computational properties of V3 and LO.

## Introduction

A central goal of vision science is to understand how the human visual system represents the shapes of objects and how shape is ultimately used to recognize objects. Research from computer vision has suggested that shape representations can be created and then compared using computational models based on the medial axis, also known as the “shape skeleton.” Although recent behavioral studies suggest that humans also represent shape skeletons (Ayzenberg & Lourenco, 2019; Firestone & Scholl, 2014), it remains unknown whether they contribute to perceptual organization, object recognition, or both. Here we provide important neural evidence that shape skeletons may be involved in both functions.

Shape skeletons are models of structure based on the medial axis of an object (Blum & Nagel, 1978). They provide a quantitative description of the spatial arrangement of object contours and component parts via internal symmetry axes (see Figure 1). Computer vision research has shown that such a description can be used to determine an object’s shape from noisy or incomplete contour information (Feldman & Singh, 2006; Kimia, 2003; Wilder et al., 2019) and to identify objects across viewpoints and category exemplars (Sebastian, Klein, & Kimia, 2004; Trinh & Kimia, 2011). Indeed, incorporating a skeletal model into off-the-shelf convolutional neural networks (CNNs) significantly improves their performance on visual perception tasks (Rezanejad et al., 2019). Similarly, behavioral research with humans has shown that participants extract the skeleton of 2D shapes (Firestone & Scholl, 2015; Kovács, Fehér, & Julesz, 1998; Psotka, 1978), even in the presence of border perturbations and illusory contours (Ayzenberg, Chen, Yousif, & Lourenco, 2019). Other research has shown that skeletal models are predictive of human object recognition (Destler, Singh, & Feldman, 2019; Lowet, Firestone, & Scholl, 2018; Wilder, Feldman, & Singh, 2011), even when controlling for other models of vision (Ayzenberg & Lourenco, 2019). Although these studies provide evidence that skeletons are associated with perceptual organization and object recognition, at least when these functions are tested independently, it remains unknown whether shape skeletons are involved in *both* functions.

**Figure 1.**
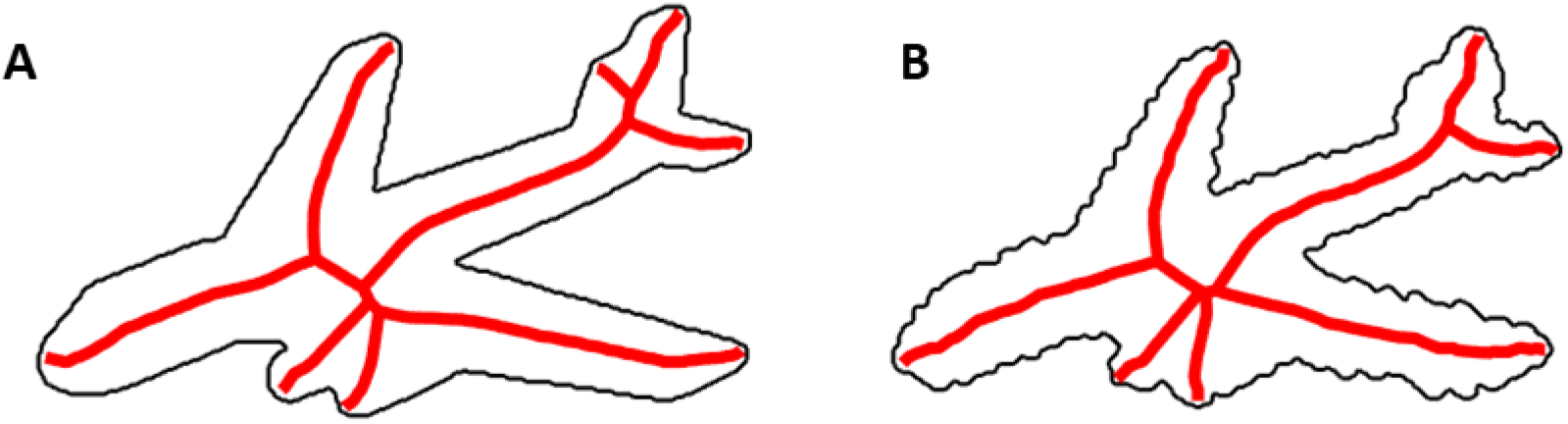
An illustration of the shape skeleton for a 2D airplane with (B) and without (A) perturbed contours. A strength of a skeletal model is that it can describe an object’s shape structure across variations in contour. Skeletons computed using the ShapeToolbox (Feldman & Singh, 2006).

One way in which to address whether shape skeletons are implicated in both perceptual organization and object recognition would be to test whether regions of the brain involved in these processes also represent the shape skeleton. If shape skeletons are used to create shape percepts, then they should be represented in areas V2, V3, and/or V4 (Ardila, Mihalas, von der Heydt, & Niebur, 2012; von der Heydt, 2015). These regions have been consistently implicated in various aspects of perceptual organization, from determining border ownership of shape contours in V2 (Zhou, Friedman, & von der Heydt, 2000) to resolving shape from motion or illusory contours in V3 and V4 (Caplovitz & Peter, 2010; Mannion, McDonald, & Clifford, 2010). If shape skeletons are also used to support object recognition, then they should also be represented in regions causally involved in recognition such as the lateral occipital cortex (LO) and/or the posterior fusiform (pFs; Freud, Culham, Plaut, & Behrmann, 2017; Grill-Spector, Kourtzi, & Kanwisher, 2001).

In one study using fMRI with humans, Lescroart and Biederman (2012) decoded skeletal structures from objects in both V3 and LO, providing preliminary support for the hypothesis that shape skeletons may be involved in both perceptual organization and object recognition. However, this study did not i) measure skeletal coding directly, leaving it unknown whether another representation of structure could account for their results, or ii) compare skeletons to other models of vision, a crucial comparison because changes to object skeletons also induce changes along other visual dimensions. Thus, the question remains about whether shape skeletons are implicated in perceptual organization, object recognition, or both.

To address this question, we created a novel set of objects that allowed us to systematically vary object skeletons and directly measure skeletal coding. We then examined the unique contributions of skeletal information to neural responses across the visual hierarchy (V1-V4, LO, pFs). More specifically, we used representational similarity analysis (RSA) to test whether a model of skeletal similarity predicted the response patterns in these regions while controlling for other models of visual similarity that approximate early-(i.e., Gabor-jet; Margalit, Biederman, Herald, Yue, & von der Malsburg, 2016), mid-(i.e., GIST and HMAX; Oliva & Torralba, 2006; Serre, Oliva, & Poggio, 2007), and high-level (i.e., AlexNet-fc6; Krizhevsky, Sutskever, & Hinton, 2012) visual processing. We also examined the robustness of skeletal representations in each region by testing whether a model of skeletal similarity generalizes across a change to the object’s component parts, which alters the non-accidental and image-level properties of the objects. Together, these comparisons and manipulations allowed us to directly measure skeletal coding in regions typically associated with perceptual organization and object recognition while ruling out alternative explanations.

## Materials and Methods

### Participants

Twenty participants (*M*_*age*_ = 19.29 years, range = 20 – 36 years; 8 females) were recruited from the Emory University community. All participants gave written informed consent to participate and had normal or corrected-to-normal vision. Experimental procedures were approved by Emory University’s Institutional Review Board (IRB). All experiments were performed in accordance with the relevant guidelines and regulations of the IRB.

### Stimuli

Twelve novel objects were selected from the stimulus set created by Ayzenberg and Lourenco (2019; see Figure 2A). The selected object set was composed of six distinct skeletons and two surface forms. The six skeletons were chosen by first conducting a *k-*means cluster analysis (*k* = 3) on skeletal similarity data for 30 unique objects (for details, see Ayzenberg & Lourenco, 2019). We selected six objects whose within- and between-cluster skeletal similarities were matched (2 per cluster). That is, the two objects from the same cluster were approximately as similar to one another as the two objects within the other clusters; objects in different clusters had comparable levels of dissimilarity to one another (see Figure 2B). This method of stimulus selection ensured that the stimulus set used in the present study contained objects with both similar and dissimilar skeletons. Two additional objects were used as targets for an orthogonal target-detection task; these objects were not included in subsequent analyses.

**Figure 2.**
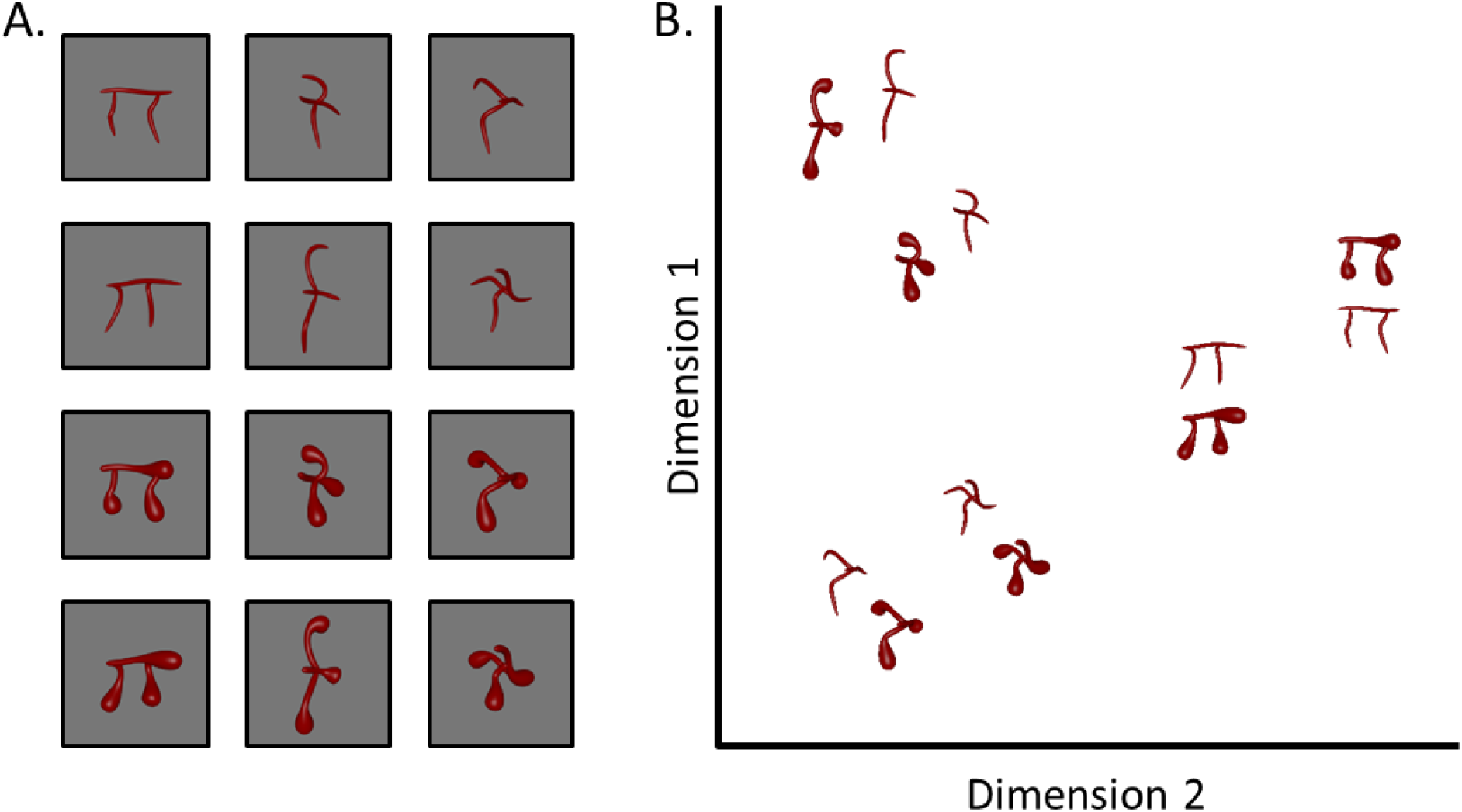
Stimuli used in the current experiment and a multi-dimensional scaling (MDS) plot illustrating the skeletal similarity between objects. (A) Six objects with unique skeletal structures were generated. Each object was rendered with two surface forms to change the objects’ component parts without disrupting the skeleton. (B) To ensure that the stimulus set contained objects with both similar and dissimilar skeletons, objects were selected in pairs such that within- and between-pair skeletal similarity were approximately matched across objects. Surface form similarity is not illustrated in the MDS plot.

Each skeleton was rendered with one of two surface forms, which changed the contours and component parts of the object without altering the underlying skeleton. To provide the strongest test of a skeletal model, we chose the two surface forms (out of five) that a separate group of participants judged to be most dissimilar (see Supplemental Materials). Importantly, we ensured that the surface forms were as discriminable as was skeletal structure (*t*(78) = 1.47, *p* = .146; see Supplemental Materials). These surface forms were also chosen because they had qualitatively different component parts, as measured by human ratings of non-accidental properties (Amir, Biederman, & Hayworth, 2012; Biederman, 1987; see Supplemental materials for details). Thus, the surface forms produced large changes to the objects as measured by participants’ discrimination judgments and ratings of non-accidental properties.

The number of objects in the current study was determined to ensure a sufficient number of presentations per object and maximize the signal-to-noise ratio. It also allowed us to implement a continuous carry-over design with third-order counterbalancing, thereby minimizing carry-over effects across trials (see Methods; Aguirre, Mattar, & Magis-Weinberg, 2011).

Finally, given that our objects were artificial and might appear to have especially salient skeletons, we tested whether they could be discriminated using visual properties other than the shape skeleton and whether they evoked visual processes similar to real objects. We found that all non-skeletal models (GBJ, GIST, HMAX, AlexNet) could accurately discriminate objects with different skeletons (80.2% - 95.3% accuracy), suggesting that skeletons were not the only available visual property and our objects differed along other visual dimensions (see Supplemental Materials for more details). Next, we tested whether participants could discriminate these stimuli in a speeded context (100 ms presentation), a task thought to evoke ‘core’ object perception (Cadieu et al., 2014; DiCarlo, Zoccolan, & Rust, 2012; see Supplemental Materials for details). This experiment revealed that participants were significantly above chance at discriminating these objects (*M* = 89.9%, *t*(13) = 14.02, *p* < .001; see Supplemental Materials). Thus, although we used artificial 3D objects designed to vary in skeletal similarity, they could also be recognized using visual properties other than the skeleton and they evoke processes typical of core object perception.

### Experimental design

First, we used a region of interest (ROI) approach, in which we independently localized the ROIs (localizer runs). Second, we used an independent set of data (experimental runs) to conduct representational similarity analyses in each ROI. Stimulus presentation was controlled by a MacBook Pro running the Psychophysics Toolbox package (Brainard, 1997) in MATLAB (MathWorks). Images were projected onto a screen and viewed through a mirror mounted on the head coil.

#### Localizer runs

We used a block design for the localizer runs. Participants viewed images of faces, bodies, objects, scenes, and scrambled objects, as previously described (Dilks, Julian, Kubilius, Spelke, & Kanwisher, 2011). Each participant completed three localizer runs, comprised of four blocks per stimulus category, each 400 s. Block order in each run was randomized. Each block contained 20 images randomly drawn from the same category. Each image was presented for 300 ms, followed by a 500 ms interstimulus interval (ISI), for a total of 16 s per block. We also included five 16 s fixation blocks: one at the beginning, three in the middle interleaved between each set of stimulus blocks, and one at the end of each run. To maintain attention, participants performed an orthogonal one-back task, responding to the repetition of an image on consecutive presentations.

#### Experimental runs

We used a continuous carry-over design for the experimental runs, wherein participants viewed images of each novel object. Each run was 360 s long. Using a de Bruijn sequence (Aguirre et al., 2011), we applied third-level counterbalancing on the image presentation order, which minimized any carry-over effects between stimuli. Importantly, this design supports smaller inter-stimulus intervals (ISIs) between stimuli (Aguirre et al., 2011; Drucker & Aguirre, 2009; Hatfield, McCloskey, & Park, 2016) and allowed for a greater number of presentations per image. Each image was presented for 600 ms, followed by a 200 ms ISI, and shown 225 times across the entire session. Each run began and ended with 6 s of fixation. To maintain attention, participants performed an orthogonal target-detection task. At the beginning of each experimental run, participants were shown one of two objects (not included in subsequent analyses) and were instructed to press a response button each time the target object appeared within the image stream.

### MRI scan parameters

Scanning was done on a 3T Siemens Trio scanner at the Facility for Education and Research in Neuroscience (FERN) at Emory University. Functional images were acquired using a 32-channel head matrix coil and a gradient echo single-shot echoplanar imaging sequence. Thirty slices were acquired for both localizer and experimental runs. For all runs: repetition time = 2 s; echo time = 30 ms; flip angle = 90°; voxel size = 1.8 × 1.8 × 1.8 mm with a 0.2 mm interslice gap. Slices were oriented approximately parallel to the anterior and posterior cingulate, covering the occipital and temporal lobes. Whole-brain, high-resolution T1-weighted anatomical images (repetition time = 1900 ms; echo time = 2.27 ms; inversion time = 900 ms; voxel size = 1 × 1 ×1 mm) were also acquired for each participant for registration of the functional images. Analyses of the fMRI data were conducted using FSL software (Smith et al., 2004) and custom MATLAB code.

### Data analyses

Images were skull-stripped (Smith, 2002) and registered to participants’ T1 weighted anatomical image (Jenkinson et al., 2002). Prior to statistical analyses, images were motion corrected, de-trended, and intensity normalized. Localizer, but not experimental, data were spatially smoothed (6 mm kernel). All data were fit with a general linear model consisting of covariates that were convolved with a double-gamma function to approximate the hemodynamic response function.

We defined regions V1-V4 bilaterally using probabilistic parcels (Wang, Mruczek, Arcaro, & Kastner, 2014). Each parcel was registered from MNI standard space to participants’ individual anatomical space. As mentioned previously, these regions were selected to test whether shape skeletons are represented in regions typically implicated in perceptual organization.

We also functionally defined object-selective region LO, as well as pFs, bilaterally in each individual as the voxels that responded more to images of intact objects than scrambled objects (*p* < 10^−4^, uncorrected; Grill-Spector et al., 1998). These regions were selected to test whether shape skeletons are represented in regions typically implicated in object recognition. Furthermore, to test the specificity of skeletal representations in object-selective regions, rather than higher-level visual regions more generally, we also defined the extrastriate body area (EBA; Downing, Jiang, Shuman, & Kanwisher, 2001) and fusiform body area (FBA; Peelen & Downing, 2005), as the voxels that responded more to images of bodies than objects (*p* < 10^−4^, uncorrected). However, because EBA shows a high degree of overlap with LO, we subtracted any EBA voxels that overlapped with LO for each participant. The same qualitative results were found when LO and EBA were not subtracted.

Analyses were conducted using the top 2000 voxels (1.8 × 1.8 × 1.8 mm) from each ROI (in each hemisphere) when available. For regions comprised of fewer than 2000 voxels, all voxels in the ROI were used (see Figure 3). To ensure that results were not related to the size of the ROI, we also conducted our primary analyses using 100, 500, and 1000 voxels. The same qualitative results were found for all ROI sizes. For each functionally defined ROI, we selected voxels that exhibited the greatest selectivity to the category of interest from the localizer runs (e.g., the 2000 most object-selective voxels in right LO). For the probabilistically-defined ROIs, we selected voxels with the greatest probability value (e.g., the 2000 voxels most likely to describe right V1). ROIs were analyzed by combining left and right hemispheric ROIs (4000 voxels total).

**Figure 3.**
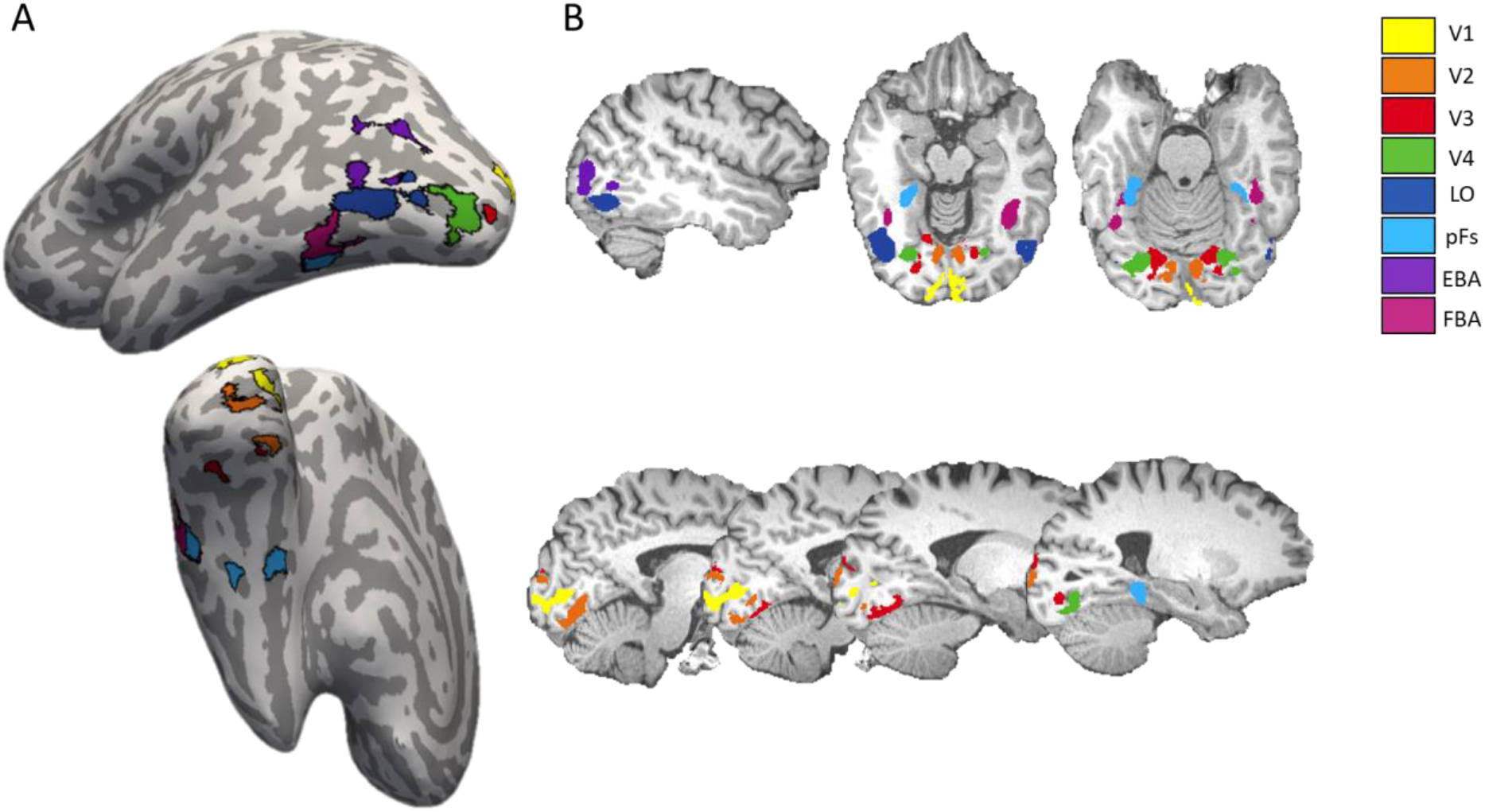
ROIs (2000 voxels) in a sample participant displayed (A) on the cortical surface and (B) in volumetric space. Each color corresponds to a different ROI. Early visual cortex ROIs (V1-V4) were defined using probabilistic maps. Higher-level visual regions (LO, pFs, EBA, FBA) were functionally defined in each participant using an independent localizer.

In subsequent analyses, we used RSA to investigate the extent to which a model of skeletal similarity explained unique variance in each ROI (Kriegeskorte, Mur, & Bandettini, 2008). For each participant, parameter estimates for each stimulus (relative to fixation) were extracted for each voxel in an ROI. Responses to the stimuli in each voxel were then normalized by subtracting the mean response across all stimuli. A 12 × 12 symmetric neural representational dissimilarity matrix (RDM) was created for each ROI and participant by correlating (1-Pearson correlation) the voxel-wise responses for each stimulus with every other stimulus in a pairwise fashion. Neural RDMs were then Fisher transformed and averaged across participants separately for each ROI. Only the upper triangle of the resulting matrix (excluding the diagonal) was used in the following analyses. Although most dissimilarity measures produce similar results, we used Pearson correlation similarity because simulations have shown it to be more reliable than other similarity measures (e.g., Euclidean distance; Walther et al., 2016).

Neural RDMs were compared to RDMs created from a model of skeletal similarity, as well as other models of visual similarity (GBJ, GIST, HMAX, and AlexNet-fc6). Skeletal similarity was calculated in 3D, object-centered, space as the mean Euclidean distance between each point on one skeleton and the closest point on the second skeleton following maximal alignment. Gabor-jet, GIST, and AlexNet (fc6-layer) similarity was calculated by extracting feature vectors from each model and computing the mean Euclidean distance between feature vectors for each feature vector. HMAX (C2-layer) similarity was calculated as the Pearson correlation between feature vectors. Because our primary analyses involve comparing the amount of unique variance explained by the skeletal model relative to the other models, we ensured that the skeletal model did not exhibit a high degree of multicollinearity with any other model, VIF = 2.61. Multicollinearity statistics for control models were also within an acceptable range (VIFs < 4.58; O’Brien, 2007).

## Results

### How are shape skeletons represented in the visual system?

We first tested whether skeletal similarity was predictive of the multivariate response pattern in each ROI by correlating the neural RDMs from each ROI with an RDM computed from a model of skeletal similarity. Significant correlations were found for V1-V4, and LO, *rs* = 0.35 – 0.67, *R*^2^ = 12.5 – 50.1 (*ps* < .001; significance determined via permutation test with 10,000 permutations; see Figure 4). Skeletal similarity was not predictive of the response pattern in pFs, EBA, or FBA (*ps* > .23), revealing specificity in the predictive power of the skeletal model (see Table 1 for correlations between the ROIs and all other models).

**Table 1.** Results of the correlation, regression, and variance partitioning analyses for each ROI and each model. Correlation analyses were conducted by correlating (1-Pearson correlation) RDMs created from the neural data from each ROI with RDMs created from each model. Regression analyses were conducted for each neural RDM by entering each model RDM as a predictor into a linear regression model. *R*^2^ values indicate the total explained variance by all of the models. Variance partitioning analyses were conducted by iteratively regressing each neural RDM on RDMs from each model and the combination of models and, then, calculating the percentage of the total explained variance (*R*^2^) uniquely explained by each model.

| ROI | Model | Model Correlations |  |  | Model Regression |  |  | Variance Partitioning |
| --- | --- | --- | --- | --- | --- | --- | --- | --- |
| | | $r$ | $r^2$ | $p$ | $R^2$ | $\beta$ | $p$ | Percentage |
| V1 |  | - | - | - | 0.52 | - | - | - |
|  | Skeleton | 0.65 | 0.43 | <.001 | - | 0.26 | .138 | 3.52 |
|  | Gabor-Jet | 0.66 | 0.44 | <.001 | - | 0.43 | .070 | 5.27 |
|  | GIST | 0.56 | 0.31 | <.001 | - | -0.09 | .618 | 0.39 |
|  | HMAX | 0.46 | 0.21 | <.001 | - | 0.14 | .297 | 1.72 |
|  | AlexNet-fc6 | 0.38 | 0.15 | .002 | - | 0.13 | .366 | 1.29 |
| V2 |  | - | - | - | 0.63 | - | - | - |
|  | Skeleton | 0.69 | 0.47 | <.001 | - | 0.23 | .140 | 2.19 |
|  | Gabor-Jet | 0.71 | 0.51 | <.001 | - | 0.55 | .010 | 7.06 |
|  | GIST | 0.60 | 0.36 | <.001 | - | -0.18 | .275 | 1.19 |
|  | HMAX | 0.53 | 0.28 | <.001 | - | 0.13 | .265 | 1.24 |
|  | AlexNet-fc6 | 0.48 | 0.23 | <.001 | - | 0.24 | .049 | 3.98 |
| V3 |  | - | - | - | 0.63 | - | - | - |
|  | Skeleton | 0.71 | 0.50 | <.001 | - | 0.46 | .003 | 8.97 |
|  | Gabor-Jet | 0.67 | 0.44 | <.001 | - | 0.10 | .637 | 0.22 |
|  | GIST | 0.62 | 0.39 | <.001 | - | 0.07 | .666 | 0.18 |
|  | HMAX | 0.56 | 0.32 | <.001 | - | 0.14 | .239 | 1.36 |
|  | AlexNet-fc6 | 0.51 | 0.26 | <.001 | - | 0.23 | .056 | 3.63 |
| V4 |  | - | - | - | 0.51 | - | - | - |
|  | Skeleton | 0.56 | 0.32 | <.001 | - | 0.18 | .321 | 1.58 |
|  | Gabor-Jet | 0.60 | 0.36 | <.001 | - | 0.41 | .085 | 4.83 |
|  | GIST | 0.55 | 0.30 | <.001 | - | -0.11 | .558 | 0.55 |
|  | HMAX | 0.50 | 0.25 | <.001 | - | 0.06 | .678 | 0.27 |
|  | AlexNet-fc6 | 0.54 | 0.29 | <.001 | - | 0.38 | .009 | 11.53 |
| LO |  | - | - | - | 0.25 | - | - | - |
|  | Skeleton | 0.35 | 0.13 | .002 | - | 0.49 | .029 | 25.46 |
|  | Gabor-Jet | 0.25 | 0.06 | .022 | - | -0.13 | .644 | 1.10 |
|  | GIST | 0.23 | 0.05 | .036 | - | -0.16 | .498 | 2.37 |
|  | HMAX | 0.33 | 0.11 | .004 | - | -0.05 | .776 | 0.42 |
|  | AlexNet-fc6 | 0.39 | 0.15 | <.001 | - | 0.41 | .020 | 29.27 |
| pFs |  | - | - | - | 0.04 | - | - | - |
|  | Skeleton | 0.07 | 0.00 | .294 | - | 0.29 | .240 | 52.33 |
|  | Gabor-Jet | -0.02 | 0.00 | .563 | - | -0.08 | .799 | 2.44 |
|  | GIST | -0.09 | 0.01 | .761 | - | -0.17 | .505 | 16.71 |
|  | HMAX | -0.07 | 0.01 | .727 | - | -0.10 | .590 | 10.90 |
|  | AlexNet-fc6 | -0.10 | 0.01 | .782 | - | 0.01 | .959 | 0.10 |
| EBA |  | - | - | - | 0.07 | - | - | - |
|  | Skeleton | 0.16 | 0.02 | .107 | - | 0.26 | .280 | 25.29 |
|  | Gabor-Jet | 0.07 | 0.01 | .287 | - | 0.04 | .900 | 0.34 |
|  | GIST | 0.00 | 0.00 | .515 | - | -0.30 | .233 | 30.91 |
|  | HMAX | 0.16 | 0.03 | .096 | - | 0.05 | .790 | 1.52 |
|  | AlexNet-fc6 | 0.12 | 0.02 | .159 | - | 0.15 | .428 | 13.57 |
| FBA |  | - | - | - | 0.06 | - | - | - |
|  | Skeleton | 0.00 | 0.00 | .484 | - | -0.19 | .442 | 16.68 |
|  | Gabor-Jet | 0.04 | 0.00 | .393 | - | 0.48 | .142 | 61.84 |
|  | GIST | -0.06 | 0.00 | .695 | - | -0.42 | .105 | 75.66 |
|  | HMAX | 0.09 | 0.01 | .242 | - | 0.12 | .515 | 11.93 |
|  | AlexNet-fc6 | 0.05 | 0.00 | .351 | - | 0.05 | .785 | 2.09 |

**Figure 4.**
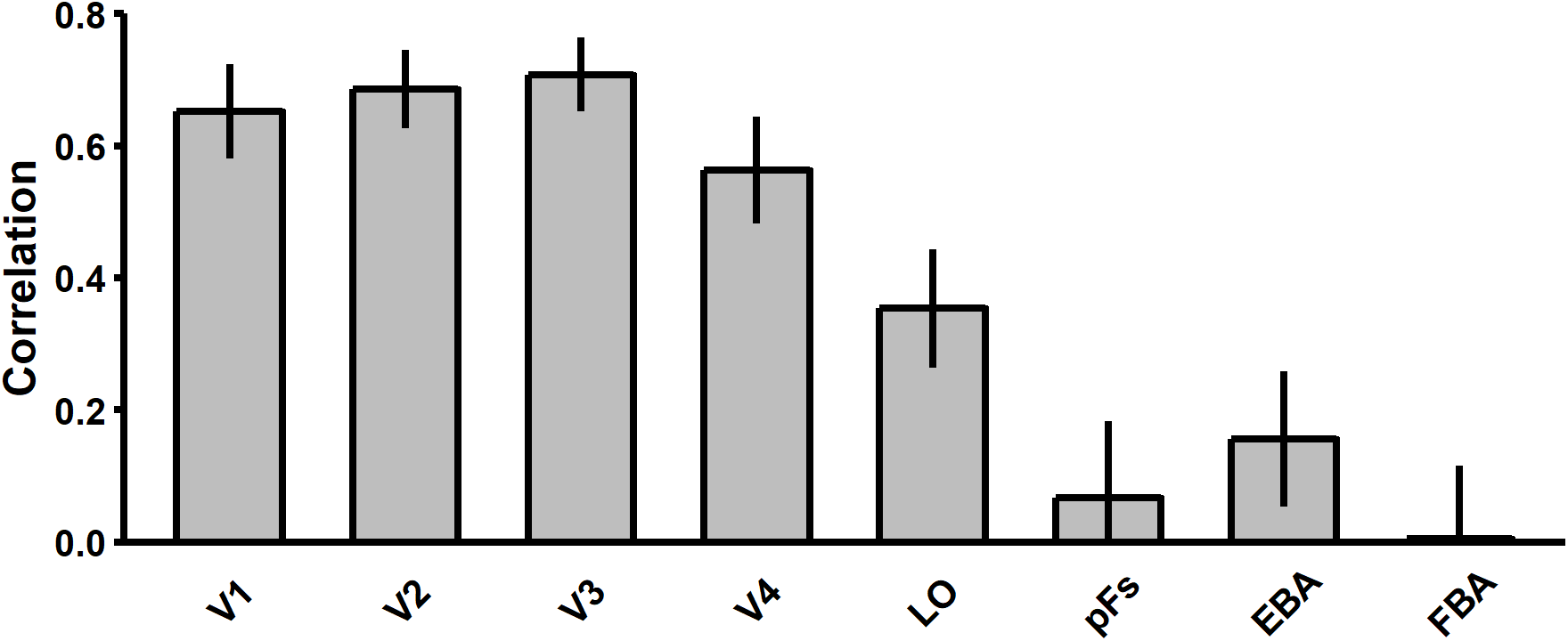
Bar plot displaying the correlations between the skeletal model and the multivariate response pattern in each ROI. A model of skeletal similarity was significantly correlated with response patterns in V1-V4 and LO. A skeletal model was not predictive of the response pattern in pFs, EBA, or FBA. Error bars represent bootstrapped SE.

Next, we tested whether skeletal similarity explained unique variance in each region or whether these effects could be explained by another model of visual similarity. To test whether the skeletal model explained unique variance in each ROI, we conducted linear regression analyses with each neural RDM as the dependent variable and the different models of visual similarity as predictors (Skeleton ∪ GBJ ∪ GIST ∪ HMAX ∪ AlexNet-fc6; see Figure 5A). These analyses revealed that the skeletal model explained unique variance in V3 (*β* = 0.46, *p* = .003) and LO (*β* = 0.49, *p* = .029), but not in the other regions (*β*s < 0.29, *p*s > .14).

**Figure 5.**
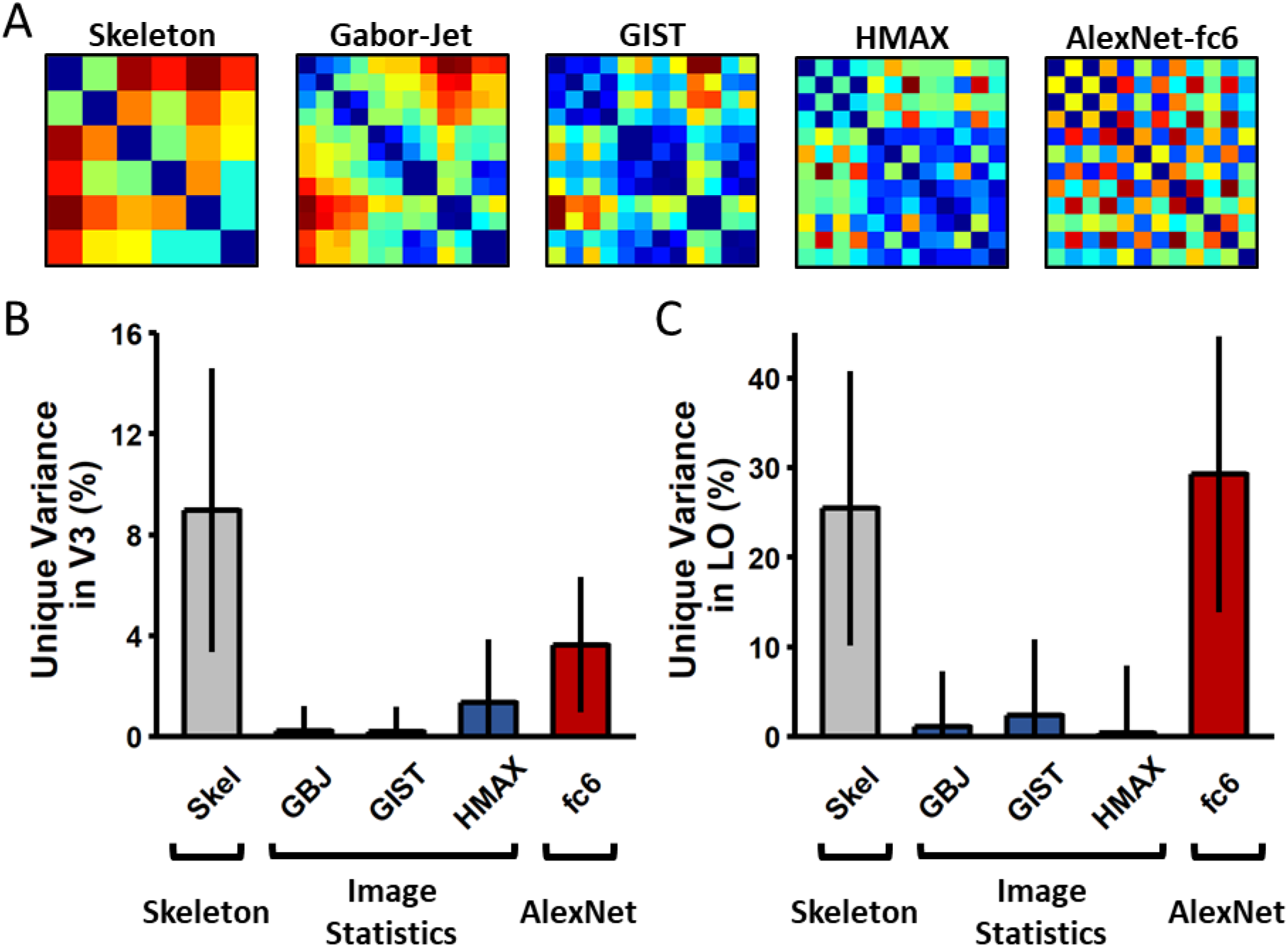
Variance partitioning results. (A) Dissimilarity matrices for the 12 stimuli computed from models of skeletal similarity and other models of visual similarity. (B) Bar plot displaying the percentage of unique variance accounted for by each model in V3. (C) Bar plot displaying the percentage of unique variance accounted for by each model in LO. A model of skeletal similarity explained unique variance in both V3 and LO, but not in other cortical regions. Error bars represent bootstrapped SE.

We also conducted variance partitioning analyses (VPA) to determine how much unique variance was explained by the skeletal model in V3 and LO (Bonner & Epstein, 2018; Lescroart, Stansbury, & Gallant, 2015). These analyses allowed us to determine how much of the total explainable variance was unique to the different models and how much was shared by a combination of models (shared variance is detailed in Supplemental Table 1). These analyses revealed that the skeletal model uniquely accounted for 9.0% of the total explainable variance in V3 and 25.5% of the explainable variance in LO (see Figure 5B-C; details about other models are provided below). Thus, shape skeletons account for significant unique variance in V3 and LO even when compared with other models of visual similarity. Together, these results are consistent with the hypothesized dual role of shape skeletons in visual processing: namely, perceptual organization and object recognition.

### Does skeletal coding in V3 and LO generalize across changes in surface form

As described previously, a strength of skeletal models is that they can be used to describe an object’s shape across variations in contours or component parts. Thus, if V3 and LO indeed incorporate a skeletal model, then these regions should represent objects by their skeletons across changes in surface form (see Figure 2). To test this prediction, new dissimilarity vectors were created from neural and model RDMs by extracting similarity values from only those object pairs whose surface forms differed and, then, correlating them to one another.

Skeletal similarity was a significant predictor of both V3 (*r =* 0.77, *p* < .001) and LO (*r* = 0.47, *p* < .001), even though object pairs were comprised of different surface forms. Notably, the finding that both V3 and LO represent shape skeletons across changes in surface form provides further evidence that skeletal coding in these regions cannot be accounted for by low-level shape properties such as contours and component parts.

But might another model of visual similarity account for these results? Here we conducted a similar regression analysis as that conducted above (neural RDMs ∼ *f*[Skeleton ∪ GBJ ∪ GIST ∪ HMAX ∪ AlexNet-fc6]) but now including subject as the random effect because fewer object pairs were involved. This analysis revealed that the skeletal model explained the greatest amount of variance in both V3 (*β* = 0.28, *p* < .001) and LO (*β* = 0.21, *p* < .001; see Supplemental Table 2 for variance explained by the other models). Thus, not only are V3 and LO sensitive to object skeletons, but these skeletal representations in these regions are also invariant to changes in surface form.

### Are V3 and LO predictive of participants’ similarity judgments of objects

Previous research has shown that shape skeletons are predictive of human participants’ behavioral judgments of object similarity (Ayzenberg & Lourenco, 2019; Destler et al., 2019; Lowet et al., 2018). Our neuroimaging results suggest that these judgments may be supported by areas V3 and LO. Here we directly tested this possibility by examining whether the response patterns of V3 and LO explain unique variance in humans’ judgments of object similarity.

Behavioral RDMs for the present objects were generated using discrimination data from Ayzenberg and Lourenco (2019). Using linear regression analyses, we first tested whether a model of skeletal similarity explained unique variance in behavioral judgments, after controlling for other models of visual similarity (GBJ, GIST, HMAX, AlexNet-fc6). We found that the skeletal model explained the greatest amount of variance in participants’ judgments (VPA = 22%, *β* = 0.28, *p* < .001), replicating Ayzenberg and Lourenco (2019). Next, we tested whether the response profiles of V3 and LO were also predictive of the behavioral RDM. These analyses revealed significant correlations for both regions and participants’ judgments: V3, *r* = 0.81, *p* < .001; LO, *r* = 0.46, *p* < .001.

In a final analysis, we tested the specificity of V3 and LO in explaining participants’ behavioral judgments by testing whether another region could explain this effect. We tested whether V3 and LO explained unique variance in participants’ judgments by conducting separate regression analyses in which the predictors were either V3 and the other early visual regions (V1, V2, V4) or LO and the other high-level visual regions (pFs, EBA, FBA). The behavioral RDM was the dependent variable in both cases. These analyses revealed that V3 (VPA = 10%, *β* = 0.83, *p* = .002) and LO (VPA = 70%, *β* = 0.70, *p* < .001) explained unique variance in participants’ similarity judgments, even when controlling for other early- and high-level visual regions, respectively.

### What role do other models of visual similarity play in the visual processing of objects

Although the skeletal model was predictive of the response profiles of V3 and LO, even across different surface forms, one might ask whether other models of visual perception still play a role in the neural processing of objects. For example, previous research has shown that other models of visual similarity account for unique variance in participants’ object similarity judgments (Ayzenberg & Lourenco, 2019; Cadieu et al., 2014; Schrimpf et al., 2018). To explore this possibility here, we tested whether these other models explained unique variance in the ROIs. Linear regression analyses revealed that the Gabor-jet model, which approximates V2-like complex cells, accounted for unique variance in the response profile of V2 (*β* = 0.55, *p* = .009), but not other regions. We also found that AlexNet-fc6, a model consisting of non-linear features, explained increasingly more variance in increasingly higher-level visual regions (V2: *β* = 0.24, *p* = .049; V4: *β* = 0.38, *p* = .009; LO: *β* = 0.41, *p* = .020). None of the models were predictive of the response profiles of V1, pFs, EBA, or FBA (*ps* > .070; see Table 1). Thus, the predictive power of these models of visual processing is largely consistent with the hypothesized regions they are meant to approximate.

## General Discussion

In the present study, we tested the hypothesis that shape skeletons are associated with two visual processes: perceptual organization and object recognition. Consistent with this hypothesis, we found that a model of skeletal similarity was predictive of the response pattern in V3, a region implicated in perceptual organization, and LO, a region involved in object recognition. Moreover, and crucially, skeletal representations in these regions could not be explained by low-, mid-, or high-level image properties, as described by other computational models of vision, nor by representations based on contours or component parts (i.e., surface forms) of the objects. These results provide novel neural evidence that the human visual system represents shape skeletons and may do so for both perceptual organization and object recognition.

The finding that V3 represents shape skeletons is consistent with human neuroimaging studies showing its involvement in perceptual organization (Sasaki, 2007). Indeed, V3 has been consistently implicated in creating shape percepts (Caplovitz, Barroso, Hsieh, & Tse, 2008; McMains & Kastner, 2010; Montaser-Kouhsari, Landy, Heeger, & Larsson, 2007) and is the earliest stage of the visual hierarchy where symmetry structure has been decoded (Keefe et al., 2018; Sasaki, Vanduffel, Knutsen, Tyler, & Tootell, 2005; Van Meel, Baeck, Gillebert, Wagemans, & Op de Beeck, 2019). But how might shape skeletons arise in V3? One possibility is that shape skeletons reflect the response profile of grouping cells (G-cells), which play an important role within neural models of perceptual organization. More specifically, these models suggest that perceptual organization is accomplished by border ownership cells (B-cells) in V2, which selectively respond to the contours of a figure (rather than the background), as well as G-cells in the subsequent visual region (e.g., V3), which coordinate the firing of B-cells via top-down connections and help to specify the contours that belong to the same figure (von der Heydt, 2015; Zhou et al., 2000). Interestingly, G-cells exhibit properties associated with shape skeletons. For example, G-cells specify the relations between contours, which may allow the visual system to determine an object’s shape despite noisy or incomplete visual information (Craft, Schütze, Niebur, & von der Heydt, 2007; Martin & von der Heydt, 2015). Moreover, the response profile of G-cells within a shape corresponds to the points of the shape’s skeleton (Craft et al., 2007), as would be expected if they implement a skeletal computation. Indeed, pruned shape skeletons, resembling those extracted from 2D shapes by human participants (Ayzenberg et al., 2019), can be generated using a model of perceptual organization that incorporates the response profile of G-cells (Ardila et al., 2012).

Nevertheless, one might ask why we did not find evidence of skeletal representation in V2 or V4, given that these regions are also frequently implicated in perceptual organization (Cox et al., 2013; McMains & Kastner, 2010; Zhou et al., 2000), particularly in electrophysiology studies with monkeys (von der Heydt, 2015). First, if shape skeletons reflect the response profile of G-cells, then they would not arise in V2, which is primarily comprised of B-cells. Moreover, G-cells are thought to arise in the visual region directly following V2 (Craft et al., 2007; Martin & von der Heydt, 2015) which, in humans, is V3 but, in monkeys, is often delineated as V4 (DiCarlo et al., 2012; Gross, Rodman, Cochin, & Colombot, 1993; Serre et al., 2007). Studies have shown that V3 in humans is proportionally much larger than in monkeys and there is debate regarding whether monkeys have a human-like V3 at all (Arcaro & Kastner, 2015; but, see Brewer, Press, Logothetis, & Wandell, 2002). Most relevant here is the fact that few studies on perceptual organization with monkeys have recorded from V3. Instead, these studies primarily focus on V2 and V4 (Hegdé & Van Essen, 2006; Poort et al., 2012; Zhou et al., 2000). Given that we found evidence of skeletal representations in V3, an intriguing possibility is that V3 may be the locus of G-cells and that skeletal representations within V3 may be an emergent property of G-cell responses.

We also found evidence of shape skeletons in LO, which is consistent with a role for skeletons in object recognition. Much work has illustrated the importance of LO in using shape for object recognition (Chouinard, Whitwell, & Goodale, 2009; Grill-Spector, Kushnir, Hendler, & Malach, 2000). This region has been shown to be particularly sensitive to object-centered shape information and is tolerant to some viewpoint changes and border perturbations (Grill-Spector et al., 2001; Grill-Spector, Kushnir, Edelman, Itzchak, & Malach, 1998). Our results suggest that LO may achieve such invariance by incorporating a skeletal description of shape, which provides a common format by which to compare shapes across variations in contours and component parts.

Importantly, our results are consistent with electrophysiology work in monkeys in which the skeletal structure of 3D objects can be decoded from monkey IT across changes in both object orientation and surface form (Hung, Carlson, & Connor, 2012). Our findings are also consistent with patient studies in which damage to LO results in a specific impairment perceiving the spatial relations of component parts, but not the parts themselves, as would be predicted by a skeletal model (Behrmann, Peterson, Moscovitch, & Suzuki, 2006; de-Wit, Kubilius, de Beeck, & Wagemans, 2013; Konen, Behrmann, Nishimura, & Kastner, 2011). Building on these studies, the present work provides the first direct evidence of skeletal representations in human LO and, crucially, demonstrates that such representations cannot be accounted for by other models of visual processing.

Interestingly, we did not find evidence of skeletal representations in another object-selective region, namely pFs. This finding may reflect a division of labor between LO and pFs, following the posterior-to-anterior anatomical gradient of shape-to-category selectivity in the ventral stream (Bracci & Op de Beeck, 2016; Freud et al., 2017). More specifically, many studies have illustrated that shape selectivity peaks in posterior regions of the ventral stream and decreases in higher-level anterior regions (Brincat & Connor, 2004, 2006; Freud et al., 2017). By contrast, sensitivity to semantic category-level information, and other non-shape visual information, progressively increases in anterior regions of the temporal lobe (Barense, Gaffan, & Graham, 2007; Behrmann, Lee, Geskin, Graham, & Barense, 2016). Given that skeletal models are exclusively descriptions of shape, such that they do not take semantic content into account, it follows that we did not find evidence of shape skeletons in pFs. Nevertheless, future research should explore this hypothesis directly by testing whether the shape skeleton explains unique variance in the response profile of pFs when familiar objects are used.

Although we found that the Gabor-jet model was predictive of the response profile in V2 (earlier in the hierarchy than shape skeletons) and that AlexNet-fc6 was most predictive in LO (same region as skeletons), the predictive value of a skeletal model in V3 and LO held even when controlling for low-(i.e., Gabor-jet), mid-(i.e., GIST, and HMAX), and high-level (i.e., AlexNet-fc6) models of visual processing. Not only are these other models representative of different levels of visual processing, but they also approximate different theories of object recognition, such as those based on image-level similarity (i.e., Gabor-jet and HMAX; Tarr & Bülthoff, 1998) and feature descriptions (i.e., AlexNet-fc6; Ullman, Assif, Fetaya, & Harari, 2016; Yamins et al., 2014). Moreover, by changing the object’s surface forms, we changed the non-accidental properties of the objects’ component parts, thereby allowing for a test of component description theories (Biederman, 1987; Kayaert, Biederman, & Vogels, 2003). These results directly address concerns of previous studies (e.g., Lescroart & Biederman, 2012) that skeletal coding could be accounted for by other models of vision and provide more direct evidence of skeletal coding in the neural processing of objects in V3 and LO.

Nevertheless, there are important caveats to the current findings. We used novel 3D objects because they allowed us to systematically control both the object skeletons and other visual features. Yet a potential concern with these stimuli is whether our results would generalize to real objects. Although we did not test this possibility directly, we predict generalizability given that participants readily discriminated these stimuli in speeded contexts, suggesting that perception of our stimuli evoke processes similar to those of real objects (DiCarlo et al., 2012). Moreover, although one might argue that such stimuli make the skeleton especially salient, not typical of other objects, we would point out that other models of vision could readily discriminate these objects and that the different surface forms directly mimic real world contexts in which different objects with the same skeletons vary in external features. Nevertheless, it is important that future work investigates the strength of these findings with real, familiar objects.

We would also acknowledge that the spatial and temporal resolution of fMRI places important qualifiers on these conclusions. First, although our results were consistent across different ROI sizes (see Supplemental Data), it is nevertheless possible that shape skeletons are represented in sub-populations of neurons within each region and that these regions have secondary functions. Indeed, V3 and LO have been shown to be sensitive to other types of visual cues, including motion (Dupont et al., 1997; Felleman & Van Essen, 1987) and depth (Parker, 2007; Welchman, 2016). Second, although we found that skeletal sensitivity peaks in V3 and LO, skeletal sensitivity may exist along a gradient within the visual hierarchy much like shape sensitivity more generally (Freud et al., 2017; Freud, Plaut, & Behrmann, 2019). Third, our data cannot address whether skeletal representations in these regions arise via feedforward or feedback processes. Indeed, feedback processes are known to be important for both perceptual organization (Mannion et al., 2010; Murray, Kersten, Olshausen, Schrater, & Woods, 2002; Wokke, Vandenbroucke, Scholte, & Lamme, 2013) and invariant object recognition (Kar, Kubilius, Schmidt, Issa, & DiCarlo, 2019; Tang et al., 2018). A more complete understanding of V3 and LO, along with experiments designed to test the causal role of shape skeletons in human vision, will be needed to confirm the claims of the present research.

In conclusion, our work highlights the unique role that shape skeletons play in the neural processing of objects. These findings not only enhance our understanding of how objects may be represented during visual processing, but they also shed light on the computations implemented in V3 and LO. Lastly, our results underscore the importance of incorporating shape information, and skeletons in particular, into models of object recognition, which currently are not implemented by most state-of-the-art CNNs (Baker, Lu, Erlikhman, & Kellman, 2018; Geirhos et al., 2018).

## Supplemental Materials

### Surface form selection

Two surface forms were used in both experiments, a ‘thin’ (Surface Form 1) and ‘bulbous’ (Surface Form 2) form. Selection of these surface forms was based on adult participants’ data from the study of Ayzenberg and Lourenco (2019). In a match-to-sample task, participants (*N* = 39) were shown one object (sample) placed centrally above two choice objects. One of the choice objects matched the sample’s skeleton, but not surface form, and the other choice object matched the sample’s surface form, but not skeleton. Participants were instructed to decide which of the two choice objects was most likely to be in the same category as the sample object. Participants performed worst at categorizing objects by their skeleton when Surface Form 1 was paired with Surface Form 2, *M* = 0.58, compared to the other surface forms (*Ms =* 0.61 - 0.78). Thus, by choosing the surface forms that presented adult participants with the greatest conflict, we provided an especially strong test of skeletal coding.

To ensure that surface forms were matched in discriminability to the selected skeletons, participants (*N* = 41) conducted a surface form discrimination task, wherein they were shown images of two objects (side-by-side) that consisted of either the same or different surface forms (same skeleton). Participants were instructed to decide whether the two images showed the same or different object. Participants discriminated between Surface Forms 1 and 2 significantly better than would be predicted by chance (0.50), *t*(40) = 8.95, *p* < .001, and importantly, discrimination accuracy between surface forms did not differ from discrimination accuracy between skeletons, *t*(80) = 0.02, *p* = .981.

In a separate set of analyses, we tested whether surface forms were comprised of qualitatively different component parts by having participants rate each surface form on the degree to which it exhibited a specific non-accidental property (NAP). During a training phase, participants (*N* = 34) were taught four NAPs (drawn from Amir et al., 2012). They then rated the degree to which each surface form exhibited a particular NAP. The four NAPs were: (1) *taper*, defined as the degree to which the thickness of an object was reduced towards the end; (2) *positive curvature*, defined as the degree to which an object part curved outwards; (3) *negative curvature*, defined as the degree to which an object part curved inwards; and (4) *convergence to vertex*, defined as the degree to which an object part ended in a point. Prior to the statistical analyses, we ensured that all participants in this sample exhibited reliable performance (αs > 0.7). A repeated measures ANOVA, with NAP as the within-subject factor and surface form as the between-subject factor, revealed a significant main effect of surface form, *F*(1, 66) = 64.00, *p* < .001, suggesting that surface forms were comprised of different NAPs.

### Can non-skeletal models discriminate the objects

A potential concern with the current study is that, because we explicitly varied skeletal similarity, object skeletons were an especially salient cue. To address this concern, we tested whether non-skeletal models could discriminate the objects used in the present study. If they could, it would suggest that the visual system need not necessarily rely on a skeletal model to discriminate these objects. A feature vector was extracted for every image (30 skeletons × 5 surface forms × 3 orientations) from each of these models (GBJ, GIST, HMAX, AlexNet-fc6). Then, for each model and object pair (same surface form), a linear support vector machine (SVM) classifier was trained to label objects using two object orientations; its ability to label the objects was tested using the third orientation. This procedure was repeated for every surface form and every combination of orientations between objects (0° × 0°; 0° × 30°; 0° × −30°; 30° × 30°; 30° × −30°; −30° × −30°). A final discrimination score was computed for each object pair by averaging the decoding accuracies across every surface form and combination of orientations. This analysis revealed that every model could discriminate between objects significantly above chance (0.50; *Ms* > 0.80), *ts* > 11.78, *ps* < .001, *ds* > 3.30 (see Supplemental Figure 1). Together, these findings demonstrate that the objects within our stimulus set were sufficiently different along other visual dimensions such that non-skeletal models could accurately discriminate them.

### Can the objects be discriminated in speeded contexts

A second potential concern is that, because we used novel stimuli, these objects did not evoke the same mechanisms as real objects, also known as ‘core’ object perception (DiCarlo et al., 2012). To rule out this possibility, we tested participants in speeded object recognition task thought to require ‘core’ object perception (Rajalingham et al., 2018). If participants maintained high accuracy on this speeded task, then it would suggest that the objects in the present study evoke the same processes as naturalistic object viewing.

Participants (*N* = 14) were administered a sequential match-to-sample task where they were asked to decide which of two choice objects matched a previously presented sample object. Each trial began with a fixation cross (500 ms), followed by a display with the sample object (100 ms), and then a display with two choice objects that remained onscreen until a response was made. One choice object had the same skeleton as the sample, whereas the other choice object had a different skeleton. The choice objects always had the same surface form as the sample (randomly selected) but were presented from different orientations (−30°, 0°, 30°). Participants were instructed to ignore the orientations of the objects and to make their decision on the basis of visual similarity. Each object was pitted against every other object an equal number of times. Each object was approximately 6° × 6° in size, and choice objects subtended 9° from the center of the screen.

Comparisons to chance (0.50) revealed that participants were able to match the sample object with the correct choice object, *M* = 89.9%, *t*(13) = 14.02, *p* < .001. This result suggests that the present stimulus set evoked processes typical of core object perception and real objects.

---

**Supplemental Figure 1.**
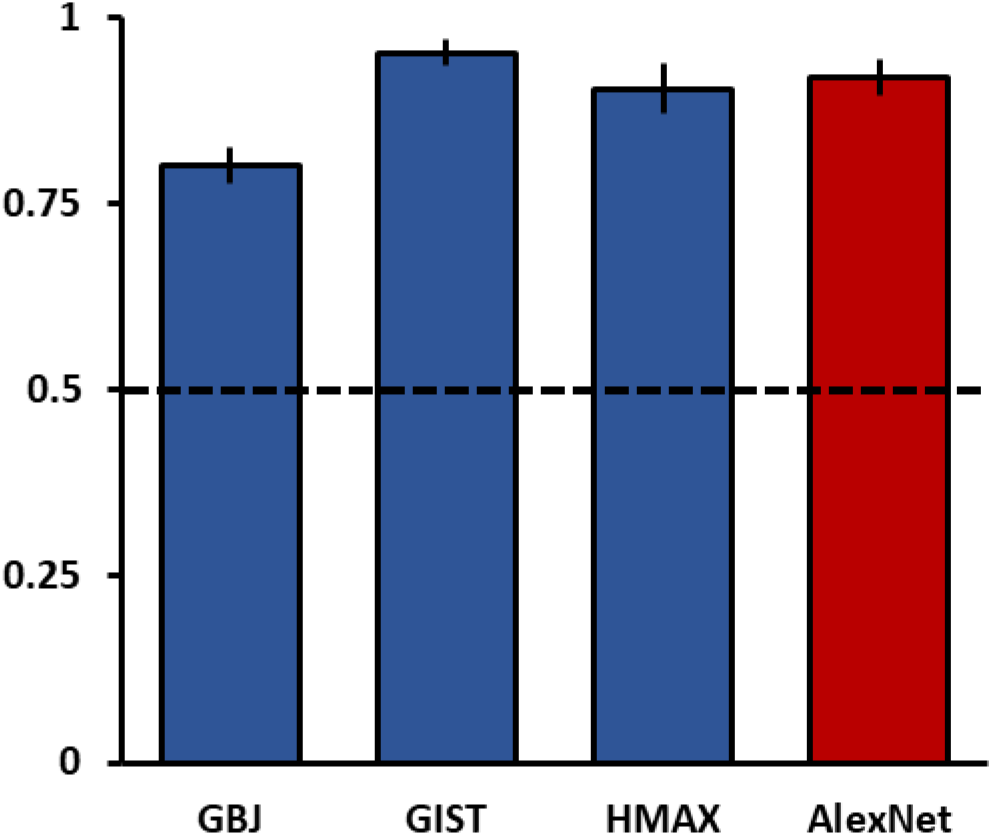
Discrimination accuracy for all non-skeletal models (proportion correct). Each model was able to discriminate between objects significantly above chance (dotted line).

**Supplemental Table 1.** Percentages of unique and shared variance explained by each model and for each ROI.

| Model | ROI |  |  |  |  |  |  |  |
| --- | --- | --- | --- | --- | --- | --- | --- | --- |
|  | V1 | V2 | V3 | V4 | LO | PFS | EBA | FBA |
| Skel | 3.5% | 2.2% | 9.0% | 1.6% | 25.5% | 52.3% | 25.3% | 16.7% |
| GBJ | 5.3% | 7.1% | 0.2% | 4.8% | 1.1% | 2.4% | 0.3% | 61.8% |
| GIST | 0.4% | 1.2% | 0.2% | 0.5% | 2.4% | 16.7% | 30.9% | 75.7% |
| HMAX | 1.7% | 1.2% | 1.4% | 0.3% | 0.4% | 10.9% | 1.5% | 11.9% |
| fc6 | 1.3% | 4.0% | 3.6% | 11.5% | 29.3% | 0.1% | 13.6% | 2.1% |
| Skel + GBJ | 18.0% | 17.5% | 11.1% | 12.2% | 9.1% | 17.8% | 28.7% | -13.9% |
| Skel + GIST | 0.3% | 0.4% | -0.2% | 0.3% | 2.3% | 8.2% | 7.7% | -7.3% |
| GBJ + GIST | 1.3% | -0.2% | 0.9% | 0.6% | 7.3% | 33.5% | 19.3% | -55.0% |
| Skel + HMAX | 2.3% | 1.5% | 3.6% | 0.7% | 0.6% | -9.7% | 7.3% | -6.7% |
| GBJ + HMAX | -0.3% | -0.3% | -0.1% | -0.1% | -0.1% | -0.6% | -0.1% | -3.1% |
| GIST + HMAX | 0.1% | 0.2% | -0.1% | 0.1% | -0.1% | -1.7% | 1.0% | 4.3% |
| Skel + fc6 | -0.6% | -0.9% | -1.7% | -1.2% | -8.4% | 0.9% | -5.6% | 2.9% |
| GBJ + fc6 | -0.3% | -0.7% | -0.1% | -1.0% | 1.0% | 0.1% | -0.2% | -1.3% |
| GIST + fc6 | -0.3% | -1.0% | 1.1% | -0.3% | -2.3% | 1.3% | -10.0% | 0.6% |
| HMAX + fc6 | 6.0% | 9.5% | 9.3% | 13.6% | 15.3% | 6.3% | 22.9% | 23.1% |
| Skel + GBJ + GIST | 28.5% | 23.2% | 24.5% | 18.3% | -11.3% | -53.3% | -48.2% | 7.2% |
| Skel + GBJ + HMAX | 6.0% | 5.4% | 4.2% | 2.8% | 2.0% | -0.6% | 9.1% | 12.8% |
| Skel + GIST + HMAX | 0.3% | 0.4% | 0.1% | 0.2% | 0.7% | 0.8% | 3.6% | 0.8% |
| GBJ + GIST + HMAX | -0.5% | -0.4% | -0.2% | -0.1% | -0.2% | -3.9% | 2.7% | 0.4% |
| Skel + GBJ + fc6 | -0.1% | -2.2% | -1.6% | -4.6% | -7.0% | 4.2% | -5.7% | -3.5% |
| Skel + GIST + fc6 | 0.0% | -0.2% | -0.8% | -0.4% | -2.2% | 7.4% | 2.4% | -4.7% |
| GBJ + GIST + fc6 | 2.1% | 3.6% | 1.6% | 6.0% | -6.5% | 5.8% | -2.6% | -0.3% |
| Skel + HMAX + fc6 | 1.2% | 1.5% | 2.7% | 1.8% | 11.1% | -2.1% | 8.4% | -4.0% |
| GBJ + HMAX + fc6 | -1.2% | -1.8% | 0.0% | -1.7% | 2.2% | -1.0% | 0.1% | -8.6% |
| GIST + HMAX + fc6 | 0.0% | -0.2% | 2.6% | 1.6% | 1.7% | 12.4% | -9.6% | -23.5% |
| Skel+ GBJ + GIST + HMAX | 4.9% | 3.8% | 4.5% | 2.4% | 0.1% | 4.9% | -10.2% | -9.3% |
| Skel+ GBJ + GIST + fc6 | -0.7% | 1.1% | 1.8% | 2.9% | 14.3% | -10.3% | 8.5% | 5.1% |
| Skel+ GBJ + HMAX + fc6 | -2.6% | -1.2% | -2.6% | 0.6% | -2.9% | 0.9% | -5.6% | 2.7% |
| Skel+ GIST + HMAX + fc6 | -0.8% | -0.7% | -0.9% | -0.6% | -2.2% | -4.7% | -6.4% | 6.0% |
| GBJ+ GIST + HMAX + fc6 | 2.5% | 3.1% | 0.6% | 2.6% | -2.3% | 7.3% | -3.6% | 11.8% |
| Skel+ GBJ + GIST + HMAX + fc6 | 21.8% | 23.1% | 25.3% | 24.7% | 19.2% | -6.5% | 14.4% | -4.5% |

**Supplemental Table 2.**
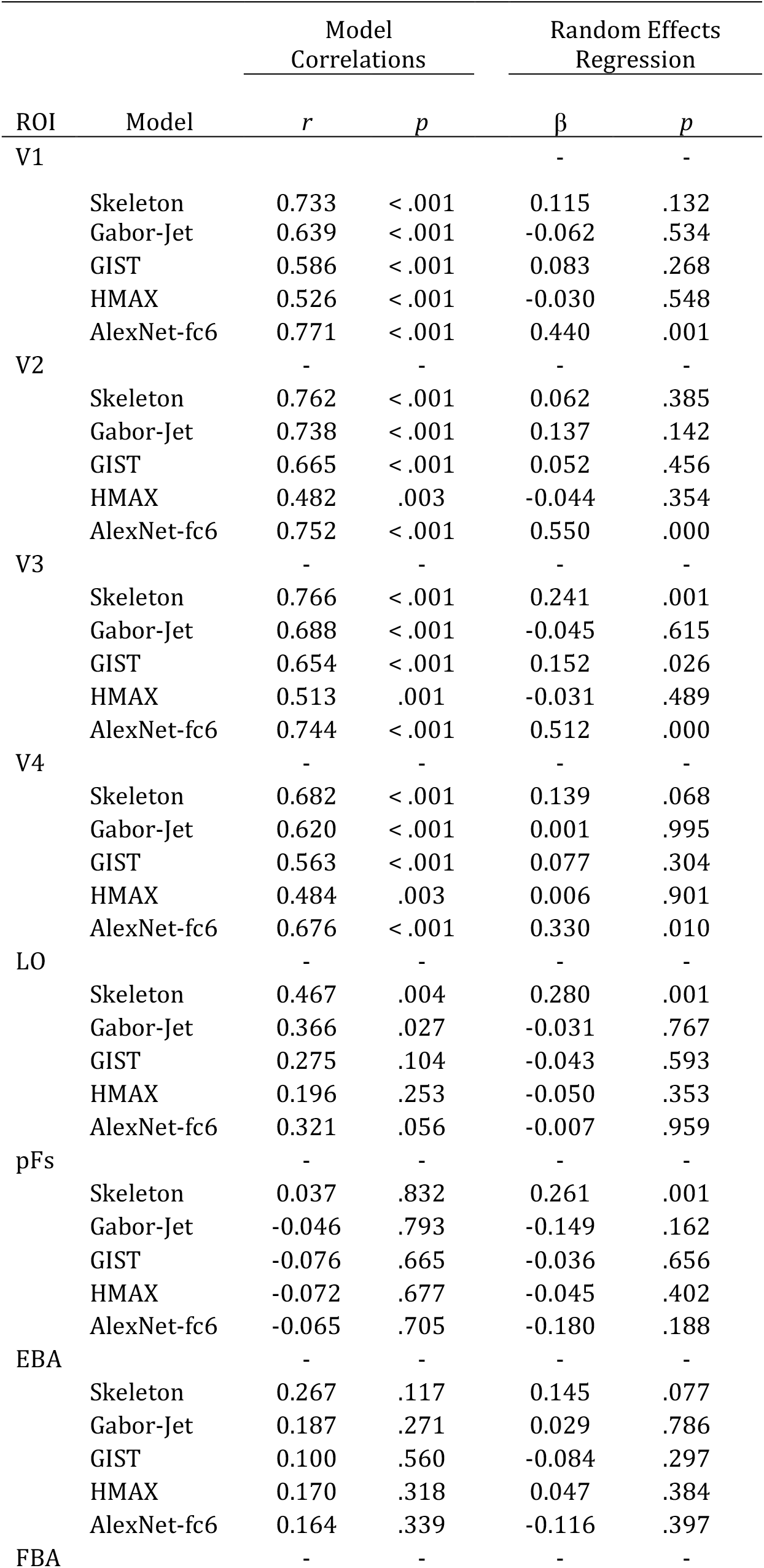
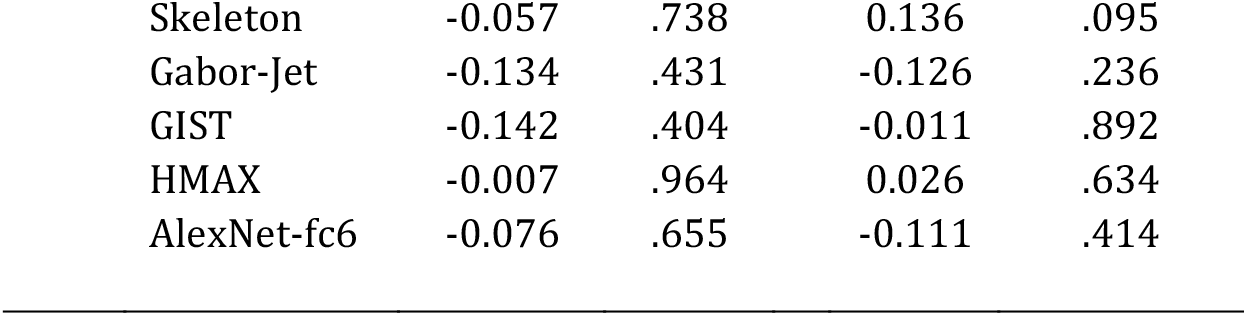
Random-effects regression results for each model and for each ROI on objects with different surface forms.

